# A potential role for reinforcement learning in speech production

**DOI:** 10.1101/2020.10.05.327072

**Authors:** Benjamin Parrell

## Abstract

Reinforcement learning, the ability to change motor behavior based on external reward, has been suggested to play a critical role in early stages of speech motor development and is widely used in clinical rehabilitation for speech motor disorders. However, no current evidence exists that demonstrates the capability of reinforcement to drive changes in human speech behavior. Speech provides a unique test of the universality of reinforcement learning across motor domains: speech is a complex, high-dimensional motor task whose goals do not specify a task to be performed in the environment but ultimately must be self-generated by each speaker such that they are understood by those around them. Across four experiments, we examine whether reinforcement learning alone is sufficient to drive changes in speech behavior and parametrically test two features known to affect reinforcement learning in reaching: how informative the reinforcement signal is as well as the availability of sensory feedback about the outcomes of one’s motor behavior. We show that learning does occur and is more likely when participants receive auditory feedback that gives an implicit target for production, even though they do not explicitly imitate that target. Contrary to results from upper limb control, masking feedback about movement outcomes has no effect on speech learning. Together, our results suggest a potential role for reinforcement learning in speech but that it likely operates differently than in other motor domains.

## Introduction

When we are speaking with someone, we are usually understood without any problems. However, sometimes this seemingly effortless communication breaks down, whether due to a noisy environment, problems in communication technology, a distracted listener, or myriad other reasons. In these situations, we need to change our speech to be better understood, but we may have limited or no information about why we were not understood or how to change our speech to maximize intelligibility. In these cases, we may try out different pronunciations of a word until we receive positive feedback from the listener that they understood what we were saying. This type of trial-by-trial learning driven by external feedback is typically often referred to as *reinforcement learning* (sometimes, as model-free learning).

Reinforcement learning has been studied extensively in upper limb control (e.g., Cashaback et al., 2017; Galea et al., 2015; Izawa & Shadmehr, 2011; Nikooyan & Ahmed, 2015; Therrien et al., 2016; Wu et al., 2014) and, to a smaller extent, in gait (Hasson et al., 2015). To date, it is essentially unknown to what extent reinforcement learning is active in speech production. Speech provides a unique test system to evaluate the universality of reinforcement learning across motor domains for two reasons. First, speech is a uniquely complex motor behavior, relying on coordination of close to roughly 100 muscles between the respiratory, phonatory, and articulatory systems that requires complex control of both skeletal joints and the tongue, a muscular hydrostat. Second, speech is unique among human motor behaviors in that the targets for movements are internally generated rather than being defined in the environment. Ultimately, the goal in speech production is to be understood, and each speaker must come to define their own motor goals to accomplish this task.

Reinforcement learning in speech may be critical both during developmental speech acquisition and for treatment of motor disorders. Developmentally, reinforcement learning has been suggested to play a critical role in the first stages of speech acquisition (Howard & Messum, 2011, 2014; Messum & Howard, 2012; Warlaumont, 2014; Warlaumont et al., 2013; Warlaumont & Finnegan, 2016). In terms of motor rehabilitation, reinforcement forms part of existing standards of care for motor speech disorders, typically combined with explicit instruction about how to produce a particular sound or set of sounds (Ballard et al., 2000; Duffy, 2013).

Despite the practical importance of reward learning in existing rehabilitation paradigms and its potential theoretical importance in human speech development, reinforcement learning in speech has received relatively little attention. The vast majority of studies on mechanisms of motor learning in speech has focused on sensorimotor adaptation–changes in behavior induced by sensory errors (e.g., Daliri & Dittman, 2019; Houde & Jordan, 1998; Lametti et al., 2012, 2018; MacDonald et al., 2011; Mitsuya et al., 2015; Purcell & Munhall, 2006; Shiller et al., 2009; Villacorta et al., 2007). While sensorimotor adaptation can drive changes in speech behavior, these changes are relatively short-lived in both speech and other motor domains compared to the longer-term impact of reinforcement learning (Krakauer, 2015; Roemmich & Bastian, 2018) and the two mechanisms rely on different neural substrates (Krakauer, 2015).

Perhaps because of its prominent role in speech rehabilitation, reinforcement learning has received some attention for clinical applications in speech (Adams et al., 2002; Adams & Page, 2000; Bislick et al., 2012, 2013; Hula et al., 2008; Katz et al., 2010; Steinhauer & Grayhack, 2000). However, these studies mostly provided highly informative feedback about performance outcomes, giving participants either explicit instruction of *how* to improve their performance or highly informative feedback about their performance such as the difference between produced duration and a duration target (often referred to as “knowledge of performance” and “knowledge of results” (Schmidt & Lee, 2011)). In non-speech domains, this type of explicit information is known to aid learning during training but often decreases retention (Hasson et al., 2015; Schmidt & Lee, 2011). Moreover, how explicit instruction interacts with other types of motor learning in unclear (Boyd & Winstein, 2004), and may in fact detrimentally affect learning in some cases (Green & Flowers, 1991; Shea et al., 2001).

The aim of the current study is to establish to what extent reinforcement learning is able to shape speech motor behavior. We additionally explore two aspects of reinforcement learning known to modulate its effectiveness in reaching. In reaching, learning is more likely when the reward signal contains some information about the desired outcome compared to uninformative signals that relay only success or failure, particularly for motor tasks involving multi-dimensional control (Kooij & Overvliet, 2016; Manley et al., 2014). Second, the presence of sensory feedback about movement outcomes has been shown to interfere with reinforcement learning in some cases (Cashaback et al., 2017).

We parametrically explore the information content of the reward signal and the availability of sensory feedback in a set of four studies on speech reinforcement learning in a 2 × 2 design. The basic goal, across all experiments, is to induce a change in the first vowel formant (F1) of the vowel /ε/ (as in *head*). Vowel formant are the characteristic resonances of the vocal tract closely tied to movements of the lips, tongue, and jaw, and are typically used to characterize vowels in speech. A similar change in vowel formants is frequently the target of sensorimotor learning studies in speech, facilitating comparison of our results with previous work in this area.

## Methods

### Participants

All participants were recruited from courses in the Linguistics and Cognitive Sciences department at the University of Delaware and were compensated with extra credit in those courses. No participant reported any history of speech or hearing problems. Experiments 1, and 2, and 4 had 20 participants each (Exp 1: 19 female/1 male; Exp 2: 20 female/0 male; Exp 4: 14 female/6 male). Experiment 3 had 21 participants (16 female/5 male). The experimental protocol was approved by Institutional Review Boards at the University of Delaware and the University of Wisconsin–Madison.

### General methods

The experiments are designed to induce participants to alter the first vowel formant (F1) in the vowel /ε/ solely through external reinforcement. Participants wore a head-mounted microphone (AKG C520) that was used to record their speech, and wore closed-back, over-the-ear headphones (Beyerdynamic DT 770) that were used to play auditory reward signals and, in Experiments 3 and 4, to play speech-shaped noise designed to mask auditory feedback. Audio data was digitized using a Scarlett 2i2 USB audio interface and recorded with the Audapter program (Cai et al., 2008; Tourville et al., 2013) in MATLAB.

Each experiment has three phases: baseline, training, and washout (Figure 1, example for “head” shown). During all phases, participants read words out loud, one at a time, as they appeared on a computer screen. Stimuli for the baseline, training, and washout phases were *head*, *bed*, and *dead* for all experiments. These stimuli contained the target vowel /ε/. Experiments 2, 3, and 4 additionally included the words *hid*, *bid*, *did* and *had*, *bad*, *dad* only during the baseline phase in order to measure F1 for the vowels /ɪ/ and /æ/, respectively. The order of the stimuli was randomized for each participant. Each word with /ε/ was repeated at least 20 times during the baseline phase of each experiment.

**Figure 1:**
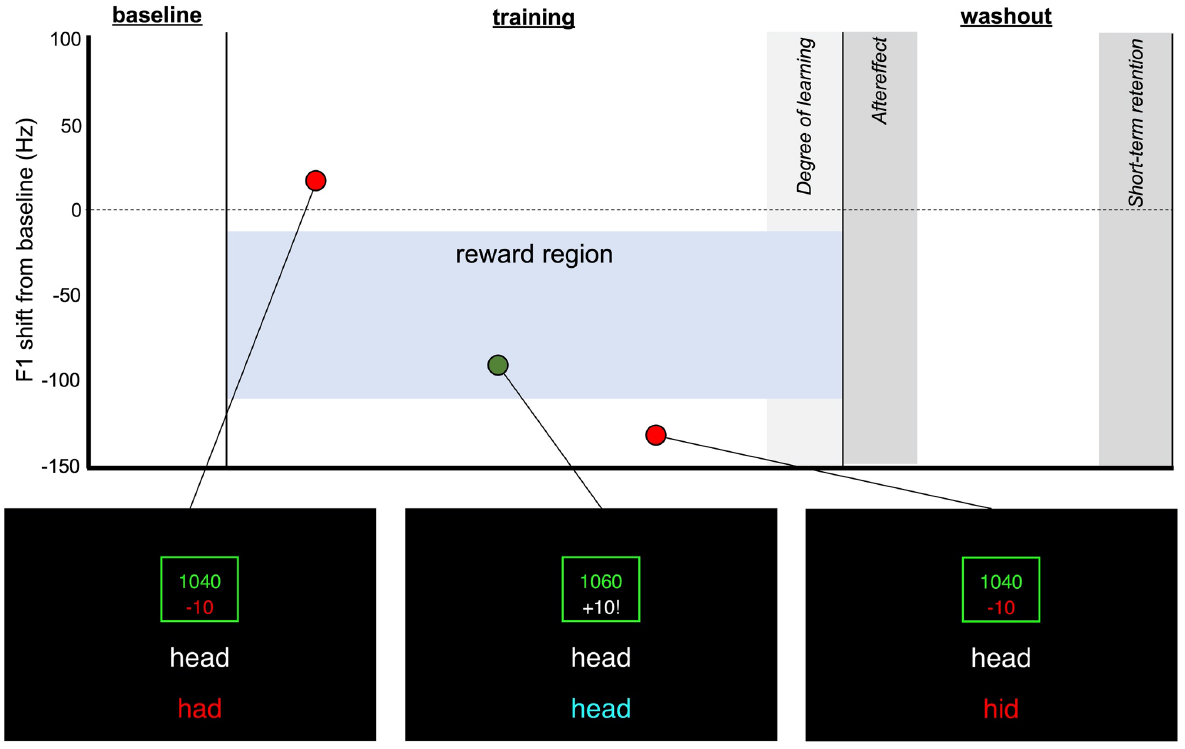
Schematic of general methods. Examples of trials with F1 above (left), in (center), and below (right) the target region are shown.

In order to provide real-time feedback based on participants’ vowel formants, the target vowel for each trial was detected automatically as the part of the speech signal above a participant-specific amplitude threshold. Formants within the detected vowel were tracked using Praat (Boersma & Weenink, 2019). A single F1 value for each trial was calculated as the average F1 within a 50 ms window centered around the vowel midpoint. Using a small window ensured the F1 measurement was taken from the steady-state portion of the vowel even with a somewhat noisy estimate of vowel onset and offset. The participant-specific amplitude threshold used for vowel detection and the Linear Predictive Coding order used for formant tracking were set in a brief parameter setting session immediately prior to the main experiment.

- <u>Baseline phase</u> (80-120 trials): Participants were told that they are training a computer program to recognize their particular voice. During this phase, the mean and standard deviation of F1 was measured. No reward or reinforcement signal was given during the baseline phase.
- <u>Training phase</u> (250-350 trials): Participants were told the computer program that was just trained will try to recognize the words they speak. Participants gained points when the computer recognized the target word (green circle in Figure 1) and lost points when it recognized another word (red circles in Figure 1). Rewards were presented visually and accompanied by auditory signals (chimes, spoken words) which varied by experiment. Participants were told that their goal was to gain points by being recognized correctly by the computer. Unknown to the participants, the computer recognized words as correct only when the first vowel formant (F1) fell within a specific target region (blue shaded region in Figure 1). This target region is 100 Hz wide, and was defined relative to the participant’s mean F1 for the vowel /ε/ produced during the baseline phase (10-110 Hz below the mean). The overlap of the reward region with participants baseline productions was chosen to ensure that participants would receive positive reward on some productions without changing their baseline behavior, as large shifts that do not overlap with baseline production may be difficult to learn (Therrien et al., 2016). A positive reward (+10 points) was given when F1 fell within a the target region. Trials with F1 values outside this region received negative reward (−10 points). Productions above the target region were recognized as containing the vowel /æ/ (e.g., *had*); those below this region, the vowel /ɪ/ (e.g., *hid*). The direction of the target region shift relative to baseline values (positive or negative) was always negative; thus, participants needed to shift their production of /ε/ towards /ɪ/ to produce F1 in the target region. Participants started with 1000 points.
- <u>Washout phase</u> (100-150 trials): Participants were told that the game is over, and that they were to simply read the words as they appear. Participants did not receive any visual or auditory feedback about the correctness of their speech or earn/lose points during the washout phase. The long washout period (100-150 trials, depending on the experiment) allowed for testing short-term retention of learning. Notably, changes in speech behavior due to sensorimotor learning return to near baseline values within 30-50 trials (MacDonald et al., 2011; Parrell et al., 2017). The washout phase was used to assess both the degree of learning (*aftereffects*, measured during first 20 trials) and short-term retention (last 20 trials).

Each trial lasted 3 seconds. Feedback about performance, if shown, was displayed for an additional 2 seconds. There was a 0.5 second pause between each trial when no stimulus word was displayed.

### Experiment-specific methods

#### Experiment 1

The baseline phase consisted of 80 trials; the training phase, 350 trials; and the washout phase, 100 trials. During the training phase, when participants’ production fell within the reward region, a pleasant chime was played over the headphones. When the production fell above or below this region, a pre-recorded voice saying the “recognized” word was played. For example, when the stimulus was “head”, “had” was played when the production was above the target region, while “hid” was played when the production fell below the target region.

#### Experiment 2

The baseline phase consisted of 120 trials; the training phase, 250 trials; and the washout phase, 150 trials. All acoustic reinforcement signals were based on each participants’ own productions recorded during the baseline phase. For each word, the production with the median F1 value in the baseline phase was chosen to be played back to the participant during the training phase. In order to create a positive reinforcement signal that fell within the target region, F1 for the chosen productions of *head, bead,* and *dead* was shifted by −60 Hz using Audapter. This resulted in an F1 in the center of the reward zone for these words. During the training, when the production fell above or below the target region, the participant’s recording of the “heard” word was played. For example, when the stimulus was “head”, “had” was played when the production was above the target region, while “hid” was played when the production fell below the reward zone. When the production fell within the reward zone, the modified version of the “heard” word was played. For example, when the stimulus was “head”, the participant’s own production of “head” from the baseline phase, with F1 shifted by −60 Hz, was played.

#### Experiments 3 and 4

Experiments 3 and 4 were designed to mirror the reinforcement signals used in Experiments 1 and 2 with the addition of speech-shaped noise designed to mask participants’ ability to hear their own speech. For Experiment 3, the baseline phase consisted of 120 trials; the training phase, 250 trials; and the washout phase, 150 trials. For Experiment 4, the baseline phase consisted of 90 trials; the training phase, 250 trials; and the washout phase, 100 trials. Stimuli with all vowels (/ɪ/, /ε/, and /æ/) were included in the baseline phase, where each stimulus word was repeated 10 times each. For both experiments, only the /ε/ stimuli were used after the baseline phase. Reinforcement signals were the same as those used in Experiment 1 (Experiment 3) and Experiment 2 (Experiment 4). The amplitude of the masking noise was modulated by the amplitude of the participant’s speech using Audapter, with the noise played at a constant gain above the speech amplitude and calibrated to be roughly 80 dB when speaking at a normal volume (Figure 2). This prevented participants from receiving auditory feedback about their speech, while largely avoiding potential Lombard affects (louder and slower speech with some changes to formant frequencies) associated with speaking in the presence of background noise (Lombard, 1911; Summers et al., 1988). A summary of differences between experiments is shown in Table 1.

**Figure 2:**
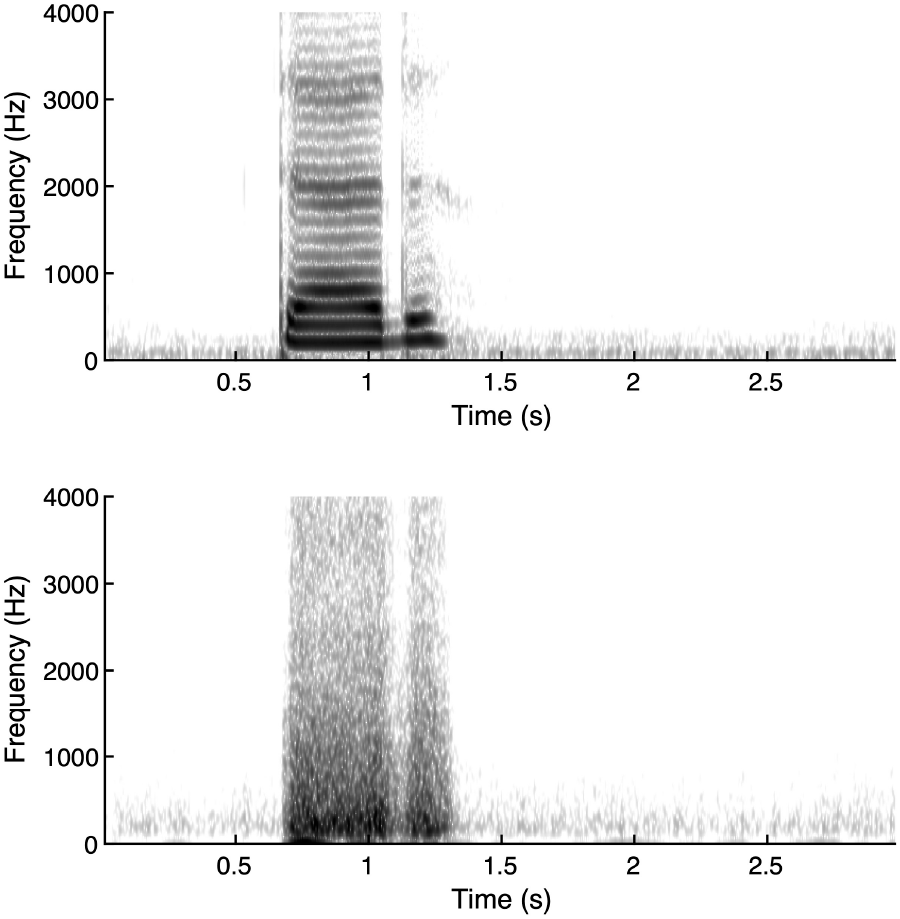
Spectrograms showing speech input (top) and amplitude-modulated masking noise (bottom) used in experiments 3 and 4. The amplitude-modulated noise served to mask auditory feedback while limiting Lombard effects associated with speaking in noise.

**Table 1:**
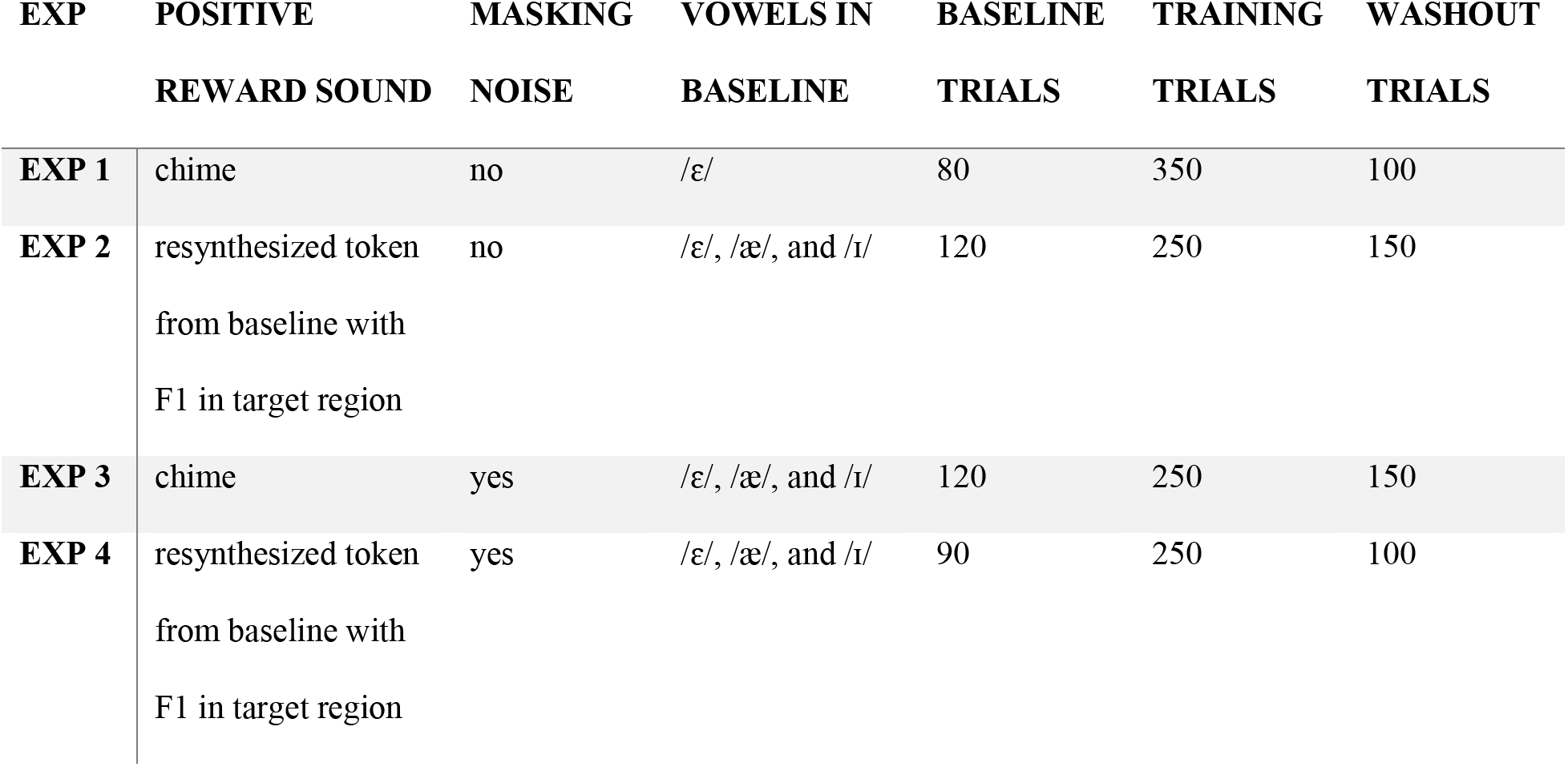
Methodological differences between experiments.

### Post-participation survey

Participants in Exp 2 and 3 were given a survey after they completed the experiment to assess whether they adopted any strategy and, if so, what that strategy was. Participants were also asked a set of questions regarding their level of engagement and attention during the experiment.

### Data analysis

The primary outcome for all experiments was the change in F1 for /ε/ from its baseline value. All trials for a given participant were normalized to the mean F1 for words with /ε/ from the baseline phase. To measure learning, we took the mean of this normalized F1 over the last 30 trials of the training phase. Aftereffects were measured as the mean F1 during the first 20 trials of the washout phase, and short-term retention was measured as the mean during the last 20 trials of the washout phase. A smaller window was chosen to measure aftereffects in order to detect potentially short-term changes in production such as those typically observed in sensorimotor adaptation studies in speech. Linear mixed-effects models were constructed using the *lme4* package (Bates et al., 2014) in *R* (R Core Team, 2013) with a fixed factor of phase (baseline, end of training, aftereffects, short-term retention) and random intercepts for participants (there were not enough observations to fit random slopes). Statistical significance was evaluated with the *lmerTest* package (Kuznetsova et al., 2017) using the Satterthwaite method to approximate the degrees of freedom. Separate tests were conducted for each experiment. Post-hoc comparisons were conducted using the *emmeans* package (Lenth et al., 2020) with Bonferroni corrections for multiple comparisons.

On visual inspection of the data, it became clear that learning was not uniform—some participants clearly showed a change speech behavior that moved their F1 to the target region, while others showed no change (Figure 3A). To quantify these differences, we sorted participants into “learners” and “nonlearners” based on their behavior in the last 30 trials of the training phase. Participants whose F1 in these trials was significantly lower than baseline (towards the target), as assessed through a t-test with α = 0.05, were classified as learners. All other participants were classified as non-learners. Classifying participants based on a metric of task success—i.e., participants who produced a significantly greater number of rewarded trials than would be expected given the standard deviation of their baseline production of words with /ε/—resulted in essentially the same classification pattern. Each method classified 2 participants as learners that were classified as non-learners by the other method. Pooling across all experiments, the distribution of learning is highly non-normal (Kolmogorov-Smirnov test: *D*(81) = 0.26, *p* < 0.0001, Figure 3B). The figure shows learning as the change in F1 from baseline to the end of the training phase, expressed as a z-score based on baseline variability. When fitting the data with two Gaussian distributions, the two distributions have centers at −2.04 and −0.15, consistent with a group of learners who lowered their F1 and a group of non-learners who did not. We report the number of learners for each experiment and descriptive statistics for learners and non-learners. However, no inferential statistics are reported for either group since the division was done a posteriori based on the data.

**Figure 3:**
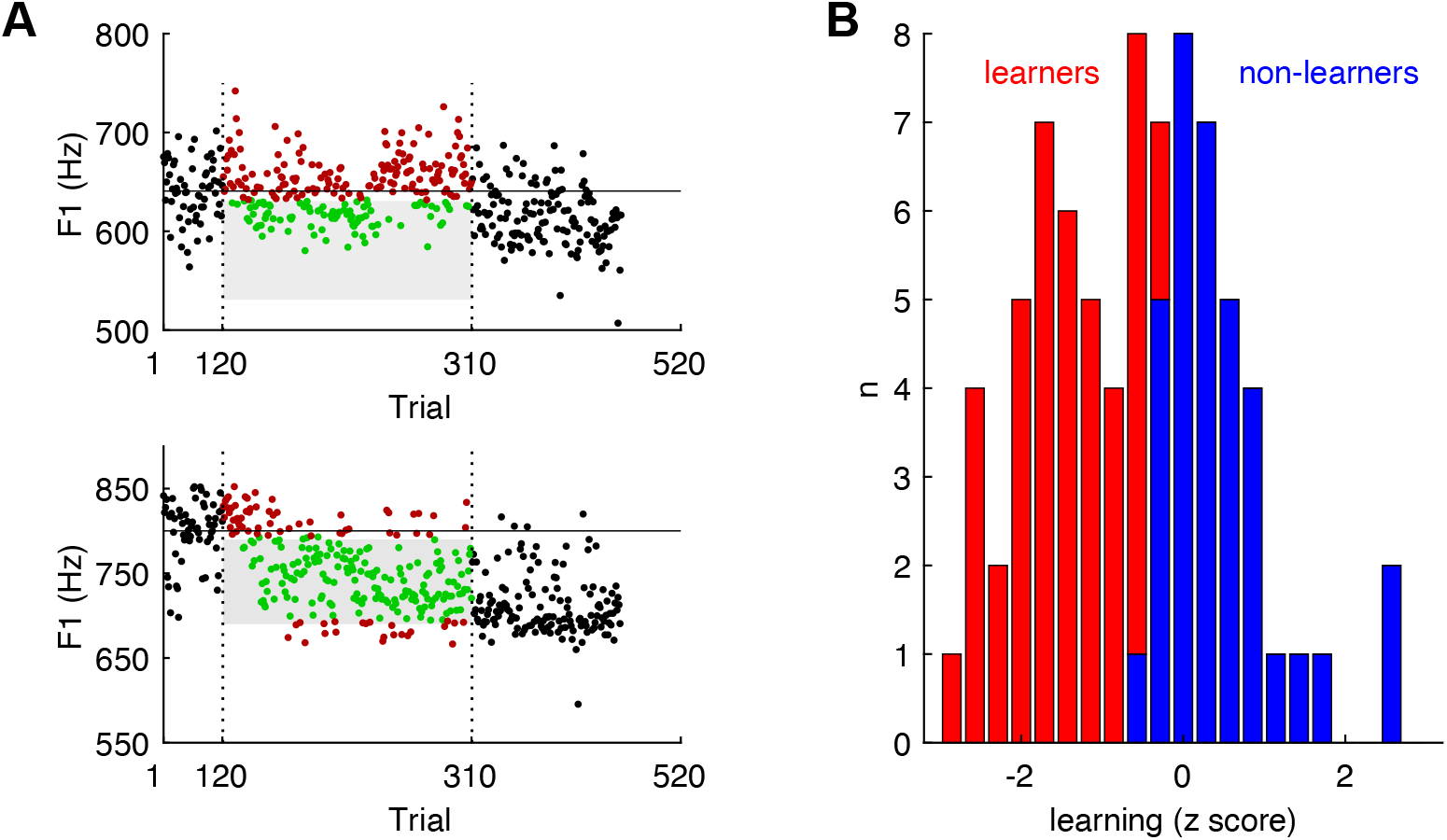
**A**: Two example participants from Experiment 3. The target region for receiving reward is shown in grey. Productions in the baseline and washout phases are shown as black circles. Productions during the training phase are shown as green circles if participants received a positive reward and as red circles if participants received a negative reward. The participant in the top panel shows no change in F1 frequency over the course of the experiment, while the participant in the bottom panel shows a clear shift in F1 frequency to the rewarded region that is maintained during washout. **B**: Distribution of learning for all participants across all four experiments. Learning shown as z-scored change in F1 from baseline values. The distribution is non-normal and has two peaks near −2 and 0. Individual participants classified as learners are shown in red; those classified as non-learners, in blue.

In addition to the individual experiment analyses, we conducted a series of cross-experiment analyses. These analyses allowed us to test directly whether the different manipulations across experiments—the type of reward signal on positively rewarded trials and the presence of masking noise—affected the degree of learning. For these analyses, we conducted an ANOVA (type 1 Sum of Squares) with reward signal and masking noise as fixed factors and the change in F1 from baseline to the end of the training phase as the dependent variable. Analyses using the aftereffects produced essentially the same results. We conducted separate analyses on both the full dataset as well as a dataset limited to only participants classified as learners, with a Bonferroni correction for multiple comparisons. This second analysis allows us to determine whether potential differences between experiments are due to different degrees of learning or, conversely, to differences in the fraction of participants who learn without any difference in the magnitude of the change in participants who do learn. To further probe whether the proportion of learners varies across experiments, we conducted Chi-squared tests comparing the proportion of learners 1) across all experiments, 2) across experiments without masking noise (Exp 1 and 2) and with masking noise (Exp 3 and 4), and 3) across experiments with no implicit imitation target (Exp 1 and 3) and with an implicit imitation target (Exp 2 and 4). A Bonferroni correction was used to correct the overall alpha level for Chi-squared tests.

A second goal of the cross-experiment analysis was to further probe the potential mechanisms driving reward learning in speech. For this, we measured another set of speech parameters related to either overall variability or trial-to-trial corrections, both of which have been suggested to be related to reward in other motor domains (Dhawale et al., 2017; Wong & Shelhamer, 2011). We measured F1 variability during the baseline phase (taken only from words with /ε/), to test whether participants who are naturally more variable may learn better. Variability was measured in two ways: as the standard deviation of all /ε/ productions in the baseline phase as well as the average trial-to-trial change in these trials. We additionally measured the change in F1 standard deviation during the first 30 training trials (early learning) compared to baseline variability to assess whether learning is associated with increased exploration of the potential solution space. We also measured the F1 distance from /ε/ to /ɪ/ during the baseline phase (Experiments 2-4 only), as participants who have a larger space between these vowels may be able to lower F1 for /ε/ without encroaching on /ɪ/. Lastly, we measured the average magnitude of the trial-to-trial change in F1 after trials with positive and negative reward. This allows us to assess how much participants change their production after a negative reward (“exploration”) and whether participants maintain similar F1 values after positive reward (“exploitation”). Statistical tests were conducted by correlating these measures with the magnitude of learning at the end of the training phase across participants, with a Bonferroni correction for multiple comparisons. Results were very similar using either aftereffects or short-term retention measures.

## Results

All experiments had the same structure (Figure 1). In all phases, participants spoke one word per trial out loud (*head*, *bed*, or *dead,* all containing the same /ε/ vowel). First, participants completed a baseline phase to measure a participant-specific mean F1 value for the vowel /ε/. No reinforcement was given during this phase. Participants were told this phase was being used to train the computer to recognize their speech. The baseline phase was followed by a training phase where participants were instructed that the computer would attempt to recognize the word they spoke, and were instructed to try to get the computer to recognize them correctly. In the training phase, the computer recognized the “correct” word if participants produced the vowel /ε/ with an F1 value 10-110 Hz below their baseline mean. Positive reward was given by earning points (+10), visual feedback of the correctly recognized word, and an experiment-specific auditory signal. In experiments 1 and 3, the auditory signal associated with positive reward was a pleasant chime. In experiments 2 and 4, the auditory signal was a token of each participant’s own speech from the baseline phase with F1 for the vowel /ε/ shifted by −60 Hz to the middle of the reward region. Negative reward was given by losing points (−10), visual feedback of the incorrectly recognized word, and an audio recording of the incorrectly recognized word. Learning was measured as the change in F1 from baseline at the end (last 30 trials) of the training phase. Following training, participants completed a washout where no reward was given. The washout phase was used to examine immediate aftereffects of learning (first 20 trials) as well as short-term retention of learning (trials 80-100). Experiments 1 and 2 had no masking noise. In Experiments 3 and 4, speech-shaped noise was played over headphones to mask participants’ ability to hear their own speech. Results for each experiment are first presented individually. All descriptive statistics show mean and standard error. Data for all experiments is shown in Figure 4.

**Figure 4:**
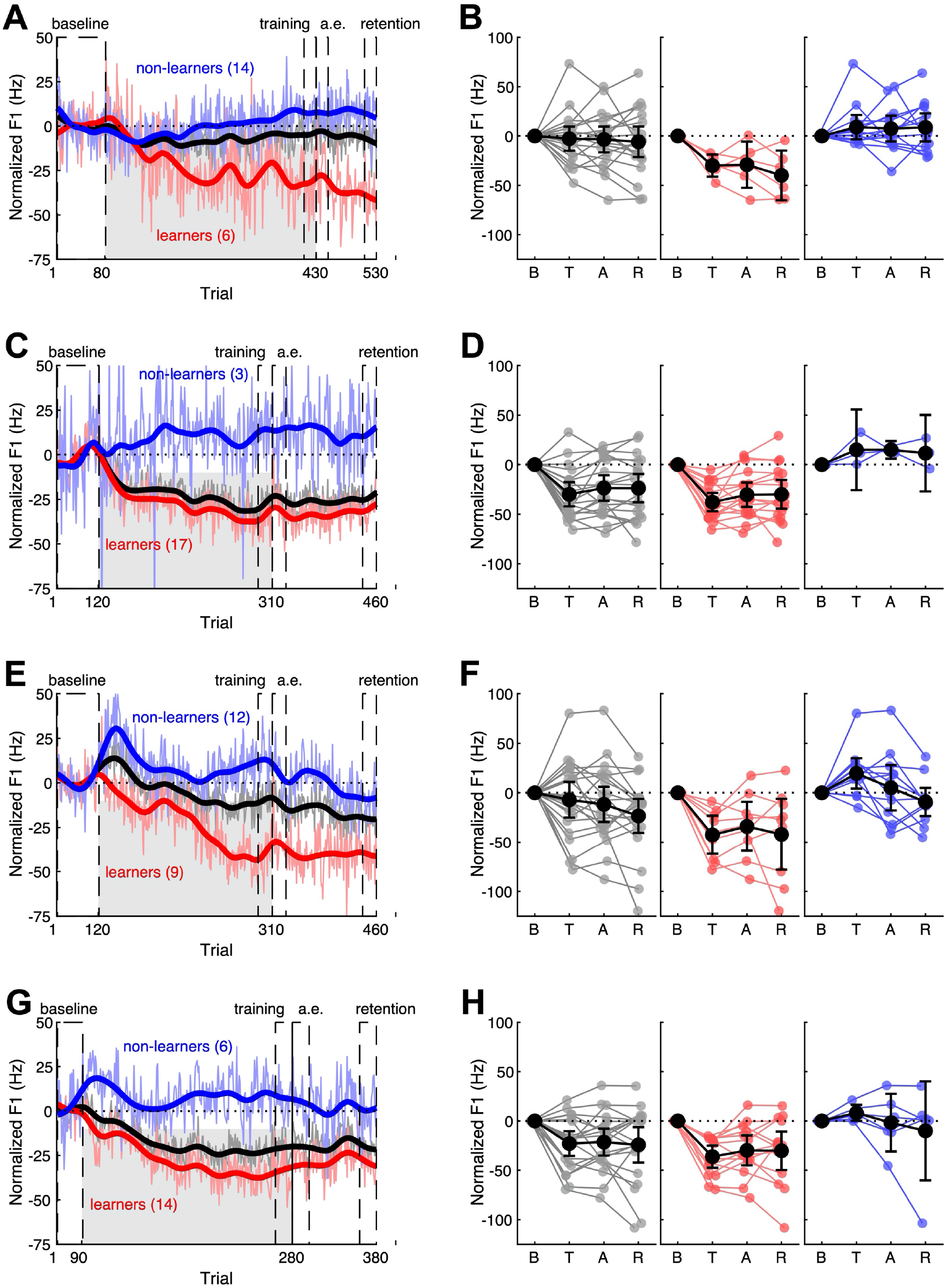
Change in F1 for all experiments. Experiments 1-4 are shown in order from the top down. **A**, **C**, **E**, **G**: mean F1 value over the course of the experiment for all participants (black), learners (red) and non-learners (blue). Raw trial averages (thin lines) as well as a smoothed running average over 10 trials (thick lines) are shown. **B, D, F, H**: F1 values in the baseline (B), end of training (T), aftereffects (A), and short-term retention (R) phases for experiments 1 (**B**), 2, (**D**), 3, (**F**), and 4 (**H**). From right to left, data is shown for all participants(back), learners (red), and non-learners (blue).

### Experiment 1

Experiment 1 had no masking noise and used a chime as the auditory signal associated with positive reward. At the group level, participants showed a very slight change in F1 values towards the target region by the end of the training phase (−2.7±5.8 Hz), which persisted into the aftereffects (−3.5±6.2 Hz) and retention (−5.9±7.4 Hz) measures. However, this change was not significant (*F*(3,57) = 0.37, *p* = 0.78). Despite the lack of an overall effect, 7/20 participants showed significant learning at an individual level, producing a change in their F1 relative to baseline values by −26.1 ± 4.8 Hz at the end of training. This change persisted into both the aftereffects (−26.6 ± 8.1 Hz) and retention (−37.1 ± 8.8 Hz) phases.

### Experiment 2

Experiment 2 had no masking noise and used a resynthesized token of each participant’s own speech, with F1 shifted to the middle of the target region as the auditory signal associated with positive reward. Participants produced a significant change from baseline after training (*F*(3,57) = 6.4, *p* < 0.0008). Across all participants, F1 was lower than baseline (*p* < 0.01) at the end of the training phase (−29.9 ± 5.8 Hz), in the aftereffects (−23.6 ± 6.1 Hz), and in retention (−23.8 ± 6.8 Hz). These phases did not differ from each other (all *p* > 0.97). At the individual level, 16/20 participants exhibited significant learning. When considering only these participants, learning was greater than for the whole group (training: −39.9 ± 4.1 Hz; aftereffects: −32.6 ± 5.7 Hz; retention: −32.2 ± 6.9 Hz).

### Experiment 3

Experiment 3 had masking noise that blocked participants’ perception of their own speech and used a chime as the auditory signal associated with positive reward. Participants did change their F1 from baseline, as reflected by a main effect of phase in the statistical model (*F*(3,60) = 3.5, *p* = 0.02). F1 was lower than baseline in all phases (training: −7.1 ± 8.6 Hz; aftereffects: −11.7 ± 8.5 Hz; retention: −23.3 ± 8.3 Hz). However, only the retention phase was significantly different from baseline (*p* = 0.01, other *p* > 0.41). The retention phase was not significantly different from either the training (*p* = 0.14) or aftereffects measures (*p* = 0.41). 9/20 participants exhibited significant learning, producing much larger changes in F1 than the group overall (training: −42.6 ± 8.4 Hz; aftereffects: −34.0 ± 10.8 Hz; retention: −42.1 ± 15.6 Hz).

### Experiment 4

Experiment 4 had masking noise that blocked participants’ perception of their own speech and used a resynthesized token of each participant’s own speech, with F1 shifted to the middle of the target region as the auditory signal associated with positive reward. Across all participants, F1 was reduced, relative to baseline, in the training (−22.9 ± 6.0 Hz), aftereffects (−21.4 ± 6.5 Hz), and retention (−24.2 ± 8.6 Hz) measures. These values were significantly lower than baseline (*F*(3,57) = 12.4, *p* < 0.0001, all individual measures *p* < 0.001). There were no differences between the three phases (all *p* > 0.63). 14/20 participants showed learning at an individual level (training: −26.2 ± 5.2 Hz; aftereffects: −34.0 ± 10.8 Hz; retention: −42.1 ± 15.6 Hz).

### Differences between experiments

In terms of overall change in F1 from baseline to the end of the training phase, there was a significant effect of auditory feedback accompanying positive reward (*F*(1,77) = 10.2, *p* = 0.002), such that the change was greater in Experiments 2 and 4, where the auditory signal was a token of each participant’s own speech with a shifted F1 value, than in Experiments 1 and 3, where the auditory signal was a chime. Contrary to our initial hypothesis, masking noise had no effect on F1 change (*F*(1,77) = 0.04, *p* = 0.85), nor was there any interaction between the presence of masking noise and the auditory signal accompanying reward (*F*(1,77) = 0.7, *p* = 0.40).

However, the effect of auditory signal type was not significant when examining only participants classified as learners (*F*(1,42) = 0.129, *p* = 0.72). Neither masking, nor the interaction between masking and reward signal were significant in this group (both *p* > 0.09). This suggests that the difference in the magnitude of F1 change between experiments with different reward signals may have been driven by differences in the proportion of learners, rather than in the degree to which participants changed F1 if they did learn. An analysis of the proportion of learners in each experiment supports this idea. There was an overall difference in the proportion of learners between all experiments (χ^2^ (3, *N* = 81) = 11.3, *p* = 0.003). This was largely driven by a difference between experiments with different reward signals (χ^2^ (3, *N* = 81) = 9.3, *p* = 0.003). There was no difference in the proportion of learners based on masking noise (χ^2^ (3, *N* = 81) = 0.3, *p* = 0.58).

Across experiments, the magnitude of F1 change at the end of the training phase was not well predicted by variability. Neither baseline variability, change in variability from the baseline to the training phase, nor distance between /ε/ and /ɪ/ in the baseline phase predicted learning (Table 2). The only predictor of learning was the trial-to-trial change in F1 after receiving positive reward. Participants who produced smaller changes in these trials learned more (*R^2^* = 0.26, *p* < 0.0001). The relationship between learning and trial-to-trial F1 changes after receiving negative reward was small and not significant after correction for multiple comparisons (*R^2^* = 0.05, *p* = 0.03). Overall, participants produced smaller magnitude F1 shifts after positive reward (32±12 Hz) than after negative rewards (41±19 Hz, t(80) = 3.6, *p* = 0.0004). Results for all factors are shown in Figure 5.

**Table 2:** Correlations between the change in F1 from the baseline to the end of the training phase and various potential predictors of learning. A * indicates a significant p-value after correction for multiple comparisons.

| MEASURE | $R^2$ | $P$ |
| --- | --- | --- |
| Baseline standard deviation | 0.01 | 0.16 |
| Baseline trial-to-trial change | -0.01 | 0.56 |
| Increase in standard error from baseline to training | -0.01 | 0.78 |
| /ε/ - /ɪ/ distance | -0.02 | 0.78 |
| F1 change after negative reward | 0.05 | 0.02 |
| F1 change after positive reward | 0.28 | < 0.0001* |

**Figure 5:**
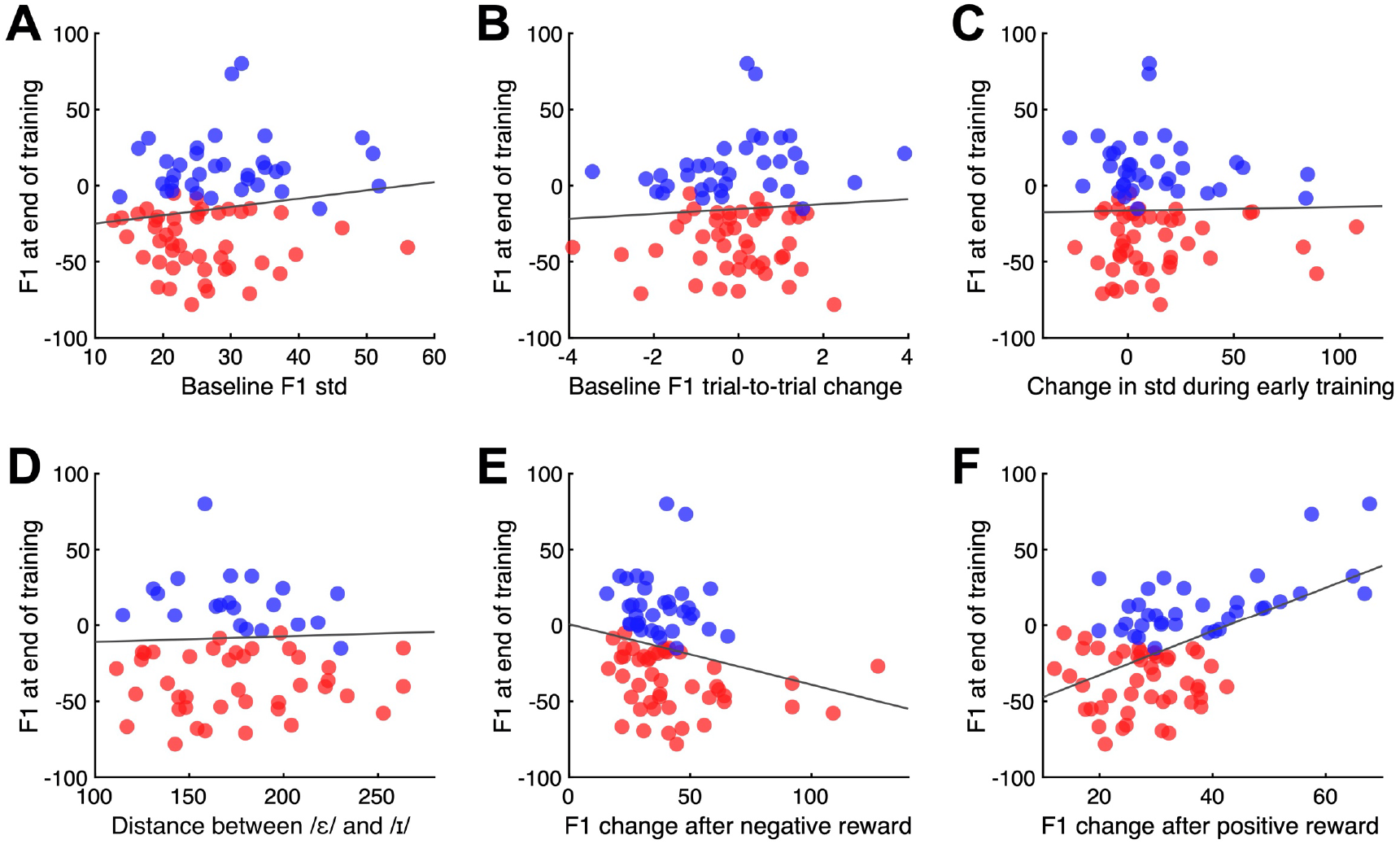
Potential factors associated with learning, defined as the magnitude of F1 change from baseline at the end of the training phase. In all panels, learners are shown in red and non-learners in blue. A solid black line indicated the regression model. **A:** Baseline standard deviation of F1. **B**: trial-to-trial change in F1 in the baseline phase. **C:** change in standard deviation of F1 from the baseline phase to the training phase. **D**: F1 distance between /ε/ and /ɪ/ in the training phase. **E**: trial-to-trial change in F1 after receiving negative reward in the training phase. **F**: trial-to-trial change in F1 after receiving positive reward in the training phase.

Based on the significant relationship between change after positive reward and learning, we considered whether the difference in overall learning magnitude (potentially driven by the proportion of learners) between experiments with informative and non-informative auditory signals accompanying reward could be related to differences in the degree to which participants shifted their productions after positive reward. For example, participants may be less likely to shift their production after they hear a word with the “correct” F1. However, we found no evidence that the magnitude of shift after positive reward differed between studies with different reward signals (*F*(1,77) = 2.4, *p* = 0.13) or based on the presence of masking noise (*F*(1,77) = 0.3, *p* = 0.59). There was similarly no significant interaction between the two factors (*F*(1,77) = 0.003, *p* = 0.96).

We additionally examined whether variability in the baseline phase or early in the training phase affected the percentage of trials that were produced with F1 in the target region. Recall that the target region ranged from 10 to 110 Hz below each participant’s baseline mean. This was chosen to ensure that all participants received reward on some trials without needing to change their baseline F1 values. Indeed, baseline variability, as measured by the standard deviation of F1, ranged from 13-56 Hz. Variability in the first 50 trials of the training phase ranged from 12-127 Hz. Even at the small end of this range, we would expect participants to receive positive reward on at least 20% of trials. In our data, all participants received at least some positive reward for trials with F1 within the target region during the training phase, as expected (1.2%-94.4% of trials, across participants). However, there was no relationship between baseline variability and percentage of trials produced with F1 in the target region across the training phase (*R^2^* = 0.03, *p* = 0.06), nor was there a relationship between variability in the training phase itself and percentage of trials with F1 in the target region (*R^2^* = 0.002, *p* = 0.774). Together, these results suggest little relationship between variability and percentage of rewarded trials.

### Strategy use and engagement

Strategy use was assessed in a follow-up survey after experiments 2 and 3. Participants were asked the question “Did you develop any techniques or strategies during the task? If so, what was that strategy?”. In Experiment 2, 16/20 participants reported using a strategy. Only 4 of these strategies related to changing the quality of the vowel, which was required to perform the task successfully. Despite the presence of a highly informative auditory signal along with positive reward (a token of the participants’ own speech with F1 shifted to the middle of the target region), only 2/20 participants reported imitating the auditory signal (both of these participants were classified as learners). In Experiment 3, 19/21 participants reported using a strategy. Of these, only 2 were plausibly related to changing vowel quality. Positive reward was accompanied by a chime in this experiment, so participants could not imitate the auditory signal. Individual participant responses are reported in the Appendix.

Participants in these studies were also asked to rate how engaging they found the task. Specifically, they were asked to rate their agreement with the statements “I was motivated to perform well in this task” and “I was motivated by the points I was earning” on a scale from 0 (disagree) to 100 (agree). The median overall motivation was 95 (mean: 84.5, 9 participants reported “yes” instead of reporting a number). The median motivation related to the points was 100 (mean: 83.5, 8 participants reported “yes” and 1 participant reported “no” instead of reporting a number).

## Discussion

In a set of four experiments, we examined whether positive and negative reinforcement could cause participants to change their speech production in the absence of any explicit instruction. Specifically, we examined whether participants could learn to lower the first formant of the vowel /ε/, analogous to a widely-demonstrated change that can be induced through sensorimotor adaptation. We tested two additional aspects of reinforcement learning. First, we examined the effects of the information content of the auditory signal associated with positive reward, comparing a non-informative sound (a chime) with a potentially-informative sound (a resynthesized version of each participant’s own speech, with F1 shifted to the center of the target region). We hypothesized that the more informative reward signal would lead to a larger magnitude of learning. Second, we examined the effect of masking auditory feedback of participants’ speech would affect learning. Based on previous work in reaching showing that visual feedback of hand position reduces the effectiveness of reinforcement learning to change reach angle, we hypothesized that learning would be reduced when auditory feedback was available, as shifting F1 in this case would conflict with participants’ internal targets for speech.

Our results provide tentative support for the ability of reinforcement to drive participants to shift their vowel production in the absence of any explicit instruction. While we observed learning in some participants in all experiments, learning at the group level was found mainly in experiments with informative auditory signals associated with positive reward that provided a reformulation of the participant’s own speech (though a small effect was also found in Experiment 3, with a non-informative auditory signal). This raises a question about whether the observed behavioral changes were driven rather by reinforcement learning *per se* or some other mechanism. It is also possible, however, that this difference in the average magnitude of learning between experiments was driven by differences in the proportion of participants who were able to learn to shift their F1 towards the target region. When examining only participants who exhibited significant learning at an individual level, the magnitude of learning was similar across studies. Thus, it may be that an informative reinforcement signal makes learning more likely, but does not affect the magnitude of learning.

It might be expected that the greater frequency of learning (and greater overall magnitude of learning) in Experiments 2 and 4 was driven by explicit imitation of the informative reinforcement signal rather than reinforcement learning. However, this appears to not be the general case. In Experiment 2, 16/20 participants were classified as learners. However, only 2/20 reported imitating the reinforcement signal. These results suggest that the benefit of an informative reward signal does not principally come from allowing for explicit imitation. However, there are a number of other possibilities for how this signal could be used that could operate separately from (or, in tandem with) reinforcement learning.

One possibility is that the reformulated feedback is being treated similarly to reafferent auditory feedback by the sensorimotor system, and that the mismatch between the produced (or predicted) formant values and formant values in the auditory feedback signal result in an auditory feedback error, analogous to those induced through altered auditory feedback paradigms (e.g., Houde & Jordan, 1998; Purcell & Munhall, 2006). If this were the case, the learning we observed in the present studies would be driven not by reinforcement but by the more well-established mechanism of sensorimotor adaptation. However, there are a number of factors which make this explanation unlikely. First, the auditory feedback played to participants had a lower F1 than the speech participants produced (on average). If this signal induced sensorimotor adaptation, it would be expected to cause an increase in F1 to oppose the perturbation; however, a *decrease* in F1 *towards* the feedback signal was observed. Second, sensorimotor adaptation in speech is highly time constrained, with even a 100ms delay in the presentation of sensory feedback drastically reducing the adaptive response, even when participants are habituated to those delays (Max & Maffett, 2015; Shiller et al., 2020). In the current experiments, shifted feedback signals were given only after the trial was over and well after the participant spoke. This is most similar to the 500ms delay condition in Max & Maffett (2015), which essentially eliminated any learning. Lastly, the effects of sensorimotor adaptation on speech are short-lived, with participants’ formant returning to near their baseline values within 30 trials. Conversely, we observed longer-lasting changes that persisted for up to 150 trials after the reward signal was removed.

A second possibility is that the resynthesized reward signal may signal how participants should change their speech, even without inducing conscious imitation. Such implicit changes could be driven by phonetic convergence or accommodation, where speakers adjust their own productions to align with speech that they hear even over very short time scales (e.g., Babel, 2010; Fowler et al., 2003; Goldinger, 1998; Pardo, 2006, 2013; Pickering & Garrod, 2013). Alternatively, this signal may have implicitly indicated the dimension of control (F1) along which speech must be altered to achieve success. Given the complex nature of speech production, this would be somewhat consistent with previous work in limb control which suggested that, in tasks with high-dimensional control, awareness of which dimensions of performance must be altered is necessary to learn from binary reinforcement (Manley et al., 2014). However, this awareness was conscious in Manley et al. (2014), while in the present case most participants did not have conscious awareness that changes in vowel quality were required.

Future work should explore to what extent varying degrees of information in the reward signal affects reinforcement learning in speech. Although the auditory signal associated with reinforcement differed across experiments, all participants received fairly informative signals associated with their performance in the form of written feedback about their production (e.g., seeing and hearing the word “had” when F1 was too high) rather than simply positive/negative feedback about performance. On the other hand, rewards given by points were binary (±10 points), and this did not carry information about how close or far participants were to the correct production. In one sense, then, the feedback given in all experiments was equivalent to signed error that indicated the direction, but not the extent, towards which speech should be changed in future repetitions.

While participants in reaching tasks may be able to use such signed error to change their aim target, it is not clear if speakers can do the same. Such “reaiming” in speech would entail conscious knowledge of 1) the acoustic relationships between different vowels in formant space (e.g., “hid” and “had” are in opposite directions from “head”) and 2) the complex transformation between speech movements and speech acoustics that would be needed to translate error in acoustic space into changes in speech movement patterns; it is unlikely naïve speakers possess such knowledge (Kim & Max, 2020). Future experiments are needed to establish precisely how speech learning is affected by participants’ knowledge of their performance as relayed both through implicit cues (e.g., graded points related to the distance to the target center instead of binary points) and cuing of the dimension of speech to be altered (e.g., explicitly informing participants they will have to change their vowel either generally or towards a specific target), as well as how these factors interact.

It will also be important to establish how more or less rewarding reinforcement signals affect learning. While participants reported to be highly motivated by the points used here, it is possible learning may be increased by more tangibly rewarding signals (e.g. monetary reward or a more immersive virtual environment). It is also possible that the reformulated speech provided a more salient or motivating feedback signal than the other sounds. The importance of speech as a highly motivating signal vs. its potential role in (implicit) imitation could be tested by using the same paradigm with *unshifted* speech tokens to accompany positive reward.

Contrary to our second hypothesis, we found no evidence that masking auditory feedback of participants’ speech affected either the magnitude or the probability of learning. This is contrary to previously demonstrated results in reaching. In these tasks, participants are presented with a visual target, and must learn to alter the angle or location of their reach away from the target to receive reward. Providing visual feedback about the position of the hand in these task seems to bias the system to weight sensory errors over reinforcement feedback, such that the effect of reinforcement on learning is eliminated (Cashaback et al., 2017). Here, we found no such effect for speech when auditory feedback is available. This may result from an important difference in how speech and reaching targets are defined. Targets in laboratory reaching tasks are externally defined (e.g., move your hand to the circle on the screen). However, movement targets in speech are defined internally by each participant. Thus, when participants change their F1 in response to reinforcement feedback, they may be simultaneously altering the intended target of their speech, eliminating any potential conflict between the sensory and reinforcement learning systems. The long-lasting aftereffects (up to 150 trials after the removal of the reinforcement signal) support the idea that participants shifted their production goals to the target region. Without anything to push them back to their pre-training targets, they maintained these goals after reinforcement was removed. More broadly, these results suggest that the interaction between sensory-error-based learning and reinforcement learning is complex and potentially reliant on whether movement targets are defined externally in the environment or internally by the individual.

Our data suggest that the primary factor driving learning is the magnitude of the change in F1 after trials that receive positive reward during the training phase. Participants who change F1 less after positive reward learn more, suggesting they are more capable of “exploiting” the correct behavior to receive reward. Perhaps surprisingly, a relationship between learning magnitude and the converse behavior of “exploring” the solution space after negative reward was not found, though participants did produce larger F1 changes after negative than positive reward overall. Learning was not related to production variability in the baseline phase or to the change in variability from the baseline to the training phase. It may have been expected that participants who were more variable were more likely to receive positive reward and thus, to learn more readily (Dhawale et al., 2017) or that higher variability in the dimension of control that must be changed would itself facilitate learning (Wu et al., 2014); however, this seems to not be the case here. This does not discount the possibility that reinforcement learning drove the observed behavioral changes, however, as other studies in reaching have similarly reported no significant relationship between learning and movement variability (Cashaback et al., 2017; Manley et al., 2014).

In sum, our results provide tentative support for the idea that reinforcement learning is an active process in speech motor control and that it can cause changes in behavior even in the absence of explicit instruction. The greatest behavioral changes were observed when reinforcement was accompanied by an informative auditory feedback signal, even though participants did not explicitly imitate that feedback. Learning was not affected by the availability of auditory feedback and learning-induced changes in production were retained after reinforcement was removed, suggesting that the learning observed here caused a shift in the intended movement target. Notably, this shift seems to be largely implicit, as few participants reported using any explicit strategies related to changing vowel quality.

These results suggest reinforcement is a plausible mechanism for early speech development, especially when coupled with “reformulations” of infant speech typically made by caregivers where they repeat the word they perceive the infant to have intended with a more adult-like pronunciation, is a plausible mechanism for early speech development (Howard & Messum, 2011; Messum & Howard, 2012; Warlaumont, 2014; Warlaumont et al., 2013; Warlaumont & Finnegan, 2016). Separately, the present results showing long-lasting behavioral changes after a relatively short training session suggest reinforcement may be a powerful clinical tool for speech rehabilitation, even without explicit instruction or detailed “knowledge of performance/results” feedback provided about errors. This has important clinical implications, as explicit instruction about how to change motor behaviors may reduce the retention of learning after training generally (Green & Flowers, 1991; Hasson et al., 2015; Shea et al., 2001; Winstein & Schmidt, 1990), and in some neurological disorders (Boyd & Winstein, 2004, 2006; Masters et al., 2004). However, potential differences between speech and other motor domains, such as the effects of sensory masking and the information content of the reward signal, suggest reinforcement learning in speech warrants further study.

## Appendix

Participant responses to post-experiment survey about strategy use.

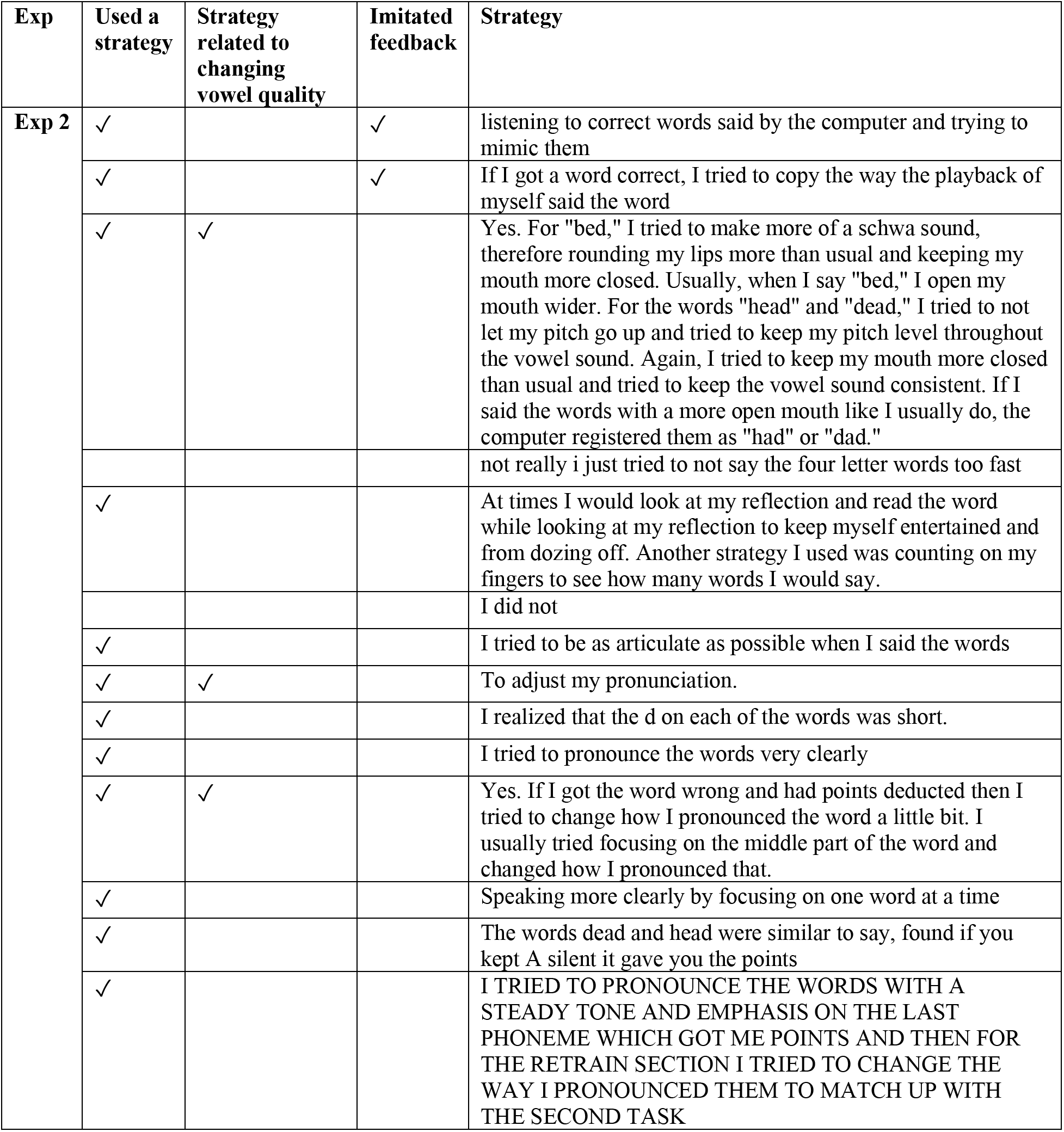
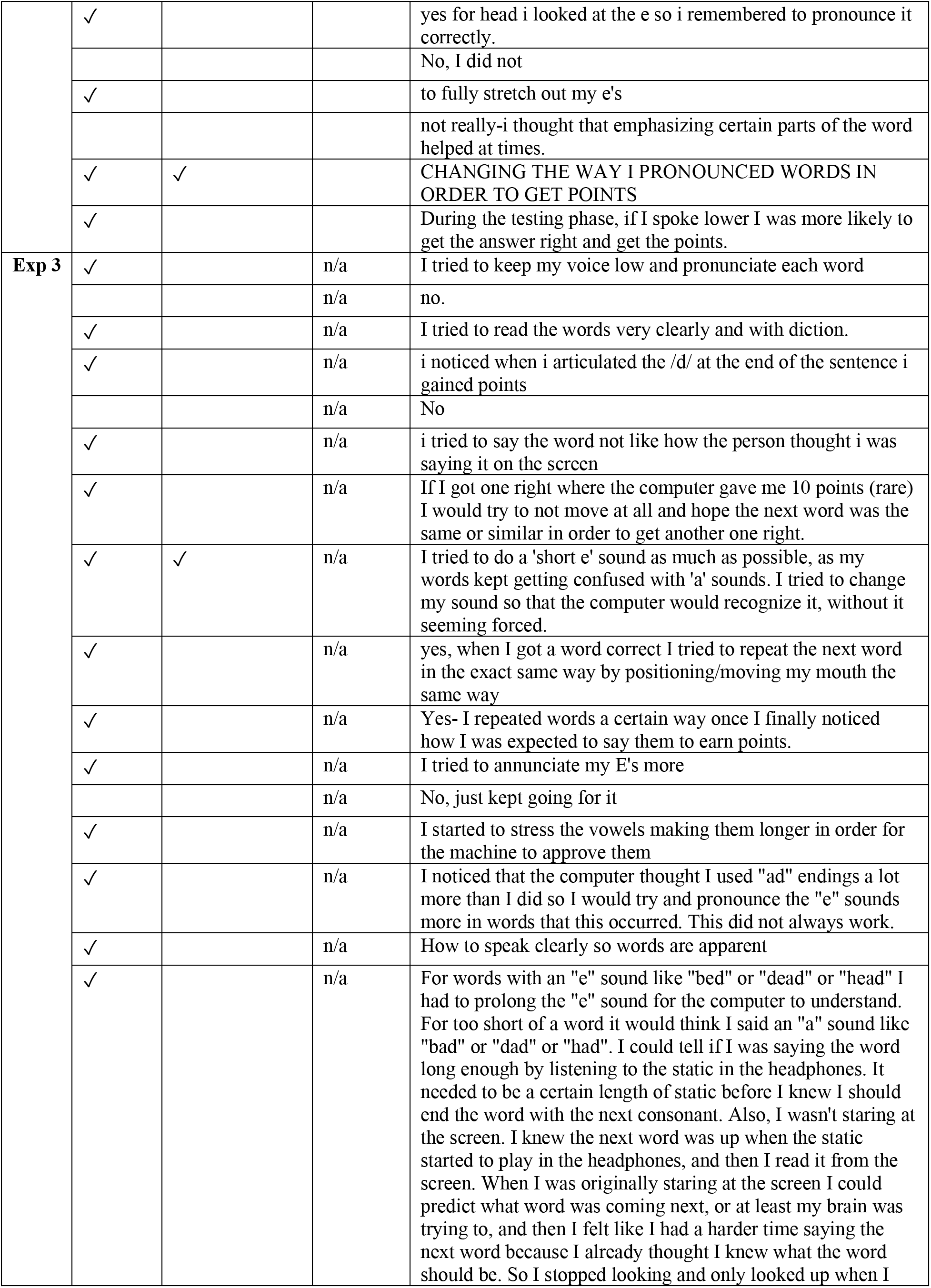
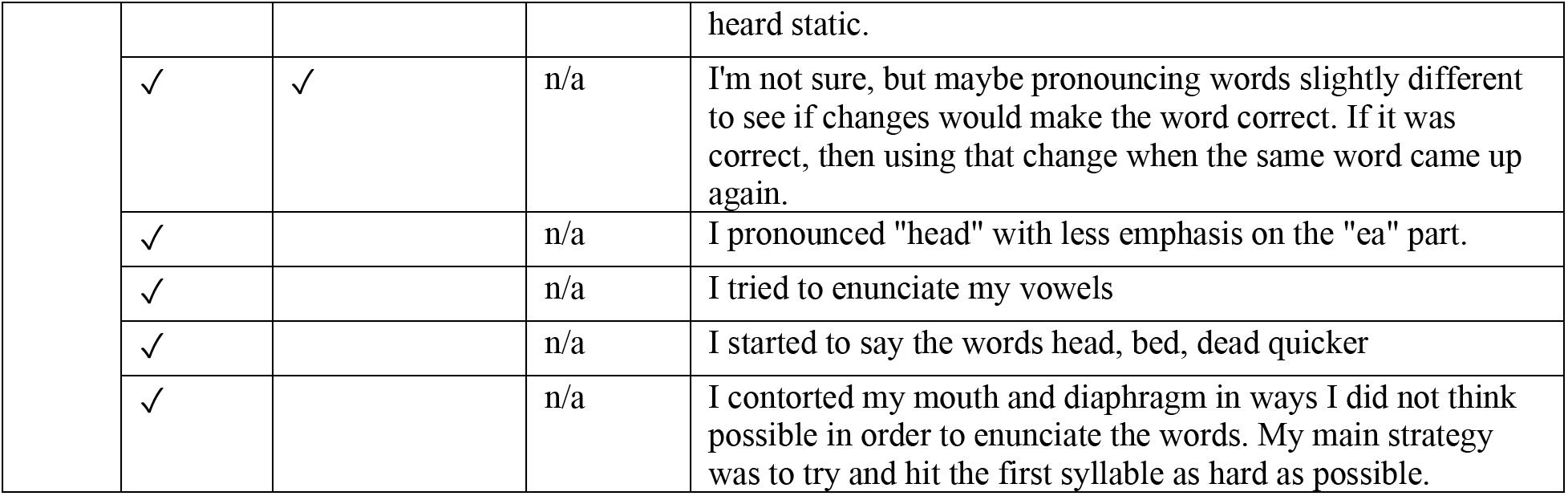

